# Prediction of early breast cancer patient survival using ensembles of hypoxia signatures

**DOI:** 10.1101/181289

**Authors:** Inna Y. Gong, Natalie S. Fox, Paul C. Boutros

## Abstract

**Background:** Biomarkers are a key component of precision medicine. However, full clinical integration of biomarkers has been met with challenges, partly attributed to analytical difficulties. It has been shown that biomarker reproducibility is susceptible to data preprocessing approaches. Here, we systematically evaluated machine-learning ensembles of preprocessing methods as a general strategy to improve biomarker performance for prediction of survival from early breast cancer.

**Results:** We risk stratified breast cancer patients into either low-risk or high-risk groups based on four published hypoxia signatures (Buffa, Winter, Hu, and Sorensen), using 24 different preprocessing approaches for microarray normalization. The 24 binary risk profiles determined for each hypoxia signature were combined using a random forest to evaluate the efficacy of a preprocessing ensemble classifier. We demonstrate that the best way of merging preprocessing methods varies from signature to signature, and that there is likely no ‘best’ preprocessing pipeline that is universal across datasets, highlighting the need to evaluate ensembles of preprocessing algorithms. Further, we developed novel signatures for each preprocessing method and the risk classifications from each were incorporated in a meta-random forest model. Interestingly, the classification of these biomarkers and its ensemble show striking consistency, demonstrating that similar intrinsic biological information are being faithfully represented. As such, these classification patterns further confirm that there is a subset of patients whose prognosis is consistently challenging to predict.

**Conclusions:** Performance of different prognostic signatures varies with pre-processing method. A simple classifier by unanimous voting of classifications is a reliable way of improving on single preprocessing methods. Future signatures will likely require integration of intrinsic and extrinsic clinico-pathological variables to better predict disease-related outcomes.

**Abbreviations:**

AUC: area under the receiver operating characteristic curve
GCRMA: GeneChip Robust Multi-array Average
HG-U133A: Affymetrix Human Genome U133A
HG-U133 Plus 2.0: Affymetrix Human Genome Plus 2.0
HR: hazard ratio
MAS5: MicroArray Suite 5.0
MBEI: Model-base Expression Index
NSCLC: Non-small cell lung cancer
RF: Random forest
ROC: receiver operator characteristic
RMA: Robust Multi-array Average

## Introduction

Cancer is fundamentally a disease driven by genetic alterations, with the stepwise accumulation of mutational hits in oncogenes and tumor suppressors [1]. However, cancer is not one disease but many, with significant variability between tumor subtypes and within individual tumours in both the rate of mutation and the specific genes that are mutated [2]. Consequently, the molecular landscape of tumours can vary wildly, leading to differences in progression and overall prognosis. These differences are described as genetic heterogeneity, while intra-tumor heterogeneity refers to heterogeneity within a tumor [3-6].

Currently, treatment decisions for individual patients are largely based on tumor subtype, histology and pathology; clinico-pathological correlation; and tumor size, nodal and metastatic status (TNM stage), along with a few molecular characteristics. This approach does not account for the wide spectrum of genetic burden experienced by the individual patients, leading to divergent responses to therapy that are currently unpredictable. Accordingly, biomarkers play a key role in the realization of precision oncology to determine the treatment that generates optimal response with minimal toxicity [7]. Biomarkers could be used at all stages of disease management, including prognosis (determining an individual patient’s likely course of disease-related outcomes such as recurrence and survival), or drug-sensitivity prediction [8, 9]. An ideal biomarker may predict multiple of these end-points simultaneously, and current research focuses on creating panels of biomarkers for each disease.

To this end, numerous groups have sought to develop transcriptomic biomarkers using microarray and RNA-sequencing approaches [7]. These efforts have resulted in a wide spectrum of signatures with prognostic potential, with the hope of fulfilling the gap between the underlying genomic heterogeneity and clinical oncology. However, few of these signatures have been successfully translated into routine clinical practice [10]. There are several reasons for this high failure rate of biomarkers [11]. First, there is little overlap in the genes incorporated across biomarkers, leading to criticism that variability in the experimental and computational techniques introduce artificial noise [12, 13]. Second, signatures have been derived from a variety of sources including cell lines, transgenic mouse models, combination of biological pathways known to be perturbed in tumor subtypes, and profiling of tumor specimens. Third, small sample size with low statistical power limits the generalizability of the signatures [14]. Fourth, biases often exist between the training and testing populations, yielding a signature that reflects interdependencies between known clinical variables [15]. Fifth, the lack of guidelines on strenuous evaluation of biomarker performance in independent validation datasets further accentuates false-positive rates and confuses the literature [14]. Finally, lack of standardized preprocessing methods challenge the consistency of the data obtained, which is often re-used in secondary studies.

Several groups have demonstrated that biomarker reproducibility is highly sensitive to the choice of preprocessing algorithm [13, 16, 17]. For example, we demonstrated that applying 24 preprocessing techniques for mRNA abundance normalization and predicting two established signatures led to only ∼33% of patients having consistent predictions in a cohort of 442 non-small cell lung cancer (NSCLC) patients [18]. Surprisingly, those patients with unanimous predictions across all preprocessing methods had more robust classifications than those from any individual preprocessing algorithm alone. These findings were corroborated when we evaluated pipeline concordance in a cohort of 1,564 early breast cancers using hypoxia signatures. The ensemble approach of merging multiple preprocessing methods improved the performance of hypoxia signatures, outperforming any individual method [19].

Hypoxia is the result of cancer altering cellular metabolism to focus on anaerobic glycolysis along with the tortuous nature of their blood vessels [19]. Hypoxic regions of the tumor have been implicated in promoting genomic heterogeneity, genomic instability and subclonal expansion of a more aggressive tumor cells [20, 21]. The selective pressures experienced by tumor cells in hypoxic conditions consequentially results in altered gene expression by epigenetics and transcription factor activation for angiogenesis, and gaining of metastatic features. Hypoxia is associated with poor prognosis and treatment failure, prompting the development of several biomarkers to identify such patients [21, 22].

It is unclear why this ensemble-of-preprocessing methods approach works so effectively. One hypothesis is that each individual preprocessing removes a different aspect of underlying noise in the microarray dataset, and that the merged ensemble of noise reduction from various perspectives allows a more accurate estimate of the true biomarker signal. The vast majority of current implementations involve simple voting, which may significantly underestimate the advantages of ensembles. Further, unanimous voting classification method leaves a large fraction (36%-80% depending on the signature) of patients unclassified. To try to bring such approaches to greater clinical utility, we set out to systematically evaluate whether ensembles of preprocessing methods may improve classification in a greater proportion of patients. We replaced the simple voting scheme with supervised machine-learning and evaluated a broad range of signatures.

## Methods

### Datasets

To systematically evaluate the impact of preprocessing ensemble classifier on risk stratification, two separate sets primary breast cancer mRNA abundance were gathered. First, eight datasets profiled on the Affymetrix Human Genome U133A (HG-U133A) microarray platform were obtained and integrated, comprising a total of 1,564 early breast cancer patients [23-30]. Second, two datasets profiled on the Affymetrix Human Genome Plus 2.0 (HG-U133 Plus 2.0) GeneChip Array were obtained for a total of 579 early breast cancer patients [31, 32]. All samples incorporated in the analysis were surgical specimens taken prior to any treatment.

### Preprocessing pipelines

To evaluate the performance of preprocessing ensemble classifiers learnt from various preprocessing pipelines, data from the two microarray platform datasets specified above were preprocessed in 24 different ways. There were three aspects that were considered to yield the unique 24 preprocessing methods: six preprocessing algorithms, two gene annotation methods, and two dataset handling procedures. The combinations of these that precipitate the 24 preprocessing pipelines were carried out as previously described [19]. Briefly, the six preprocessing algorithms include 4 without log_2_-transformation [Robust Multi-array Average (RMA) [33], MicroArray Suite 5.0 (MAS5) [34], Model-base Expression Index (MBEI) [35], GeneChip Robust Multi-array Average (GCRMA) [36]], and 2 log_2_-transform versions of MAS5 and MBEI. These algorithms were all available in the R statistical environment (R packages: affy v1.36.0, gcrma v2.30.0). **Additional file 1: Table S1** provides a brief summary of each of these algorithms. The two dataset handling approaches include either independent or merged preprocessing. The two ProbeSet annotations used were either default Affymetrix gene-annotation (R packages: hgu133aprobe v2.10.0, hgu133acdf v2.10.0, hgu133a.db v2.8.0, hgu133plus2probe v2.6.0, hgu133plus2cdf v2.6.0, hgu133plus2.db v2.8.0) or an alternative Entrez Gene-based updated annotation (R packages: hgu133ahsentrezgprobe v15.1.0, hgu133ahsentrezgcdf v15.0.0, hgu133plus2hsentrezgprobe v15.1.0, hgu133-plus2hsentrezgcdf v15.1.0). **Additional file 2: Table S2** provides a summary of each of these preprocessing pipelines.

### Patient risk classification: hypoxia signatures

To assess the influence of preprocessing variation on risk stratification of patients, we used four published hypoxia gene signatures: Buffa metagene [37], Winter metagene [38], Hu signature [39], and Sorensen gene set [40]. These signatures were chosen as they exhibited the best performance in predicting patient outcome in our previous work. Briefly, each gene signature was used to stratify patients into either low-risk or high-risk. Following pre-processing of data using pipelines, the multi-gene signature score was calculated for each patient using all genes on the signature’s gene list. First, for each gene of the signature, patients were median dichotomized (0 or 1) based on the signal-intensity of the gene compared to the expression level of that gene across all patients. Next, the multi-gene signature score for each patient was calculated as the sum of all gene scores. Finally, the scores were used to median dichotomize patients into high and low risk groups for each signature.

For preprocessing pipelines with independent dataset preprocessing, stratification was conducted independently. In preprocessing pipelines with merged dataset preprocessing, stratification was conducted simultaneously. In summary, for each patient, 24 risk classifications (high or low risk) was derived from 24 different pre-processing pipelines based on gene signature expression.

Brief descriptions of the original studies deriving these signatures are provided in **Additional file 3: Table S3**. Of note, genes contained in these signatures are genes that were found to be upregulated in hypoxic tumor environments, resulting in worse prognosis.

### Ensemble classifier: risk classification votes

The primary endpoint was to delineate whether an ensemble of preprocessing pipeline classifiers using hypoxia signatures may improve the prediction of prognosis in early breast cancer patients beyond that achieved by single pipeline classifiers. Since cause-specific mortality data is lacking in our study, individual patient survival outcome was defined as either 0 or 1 to represent dead or alive status at 5-years, respectively (events occurred after 5-years were censored). Five-year survival was chosen as it is an important survival time-point for breast cancer survivors due to the increasing causes of death unrelated to breast cancer in subsequent survivorship years. At the end of 5 years, 1193 were censored while 371 cancer-related events occurred for patients profiled on the HG-U133A platform. For patients profiled on the HG-U133 Plus 2.0 platform, 352 were censored while 227 events occurred.

The 24 dichotomized risk profiles determined from each hypoxia signature were combined to develop a preprocessing ensemble classifier using random forest (randomForest package v4.6.10) to stratify patients within the HG-U133A and HG-U133A Plus 2.0 datasets, respectively, as good or poor prognosis. The HG-U133A and HG-U133A Plus 2.0 datasets were independently separated into training and testing sets by a sample size ratio of 1:1. Random sampling was employed to determine the training and testing set, maintaining a balanced ratio between mortality and survival events in subsequent datasets. Random forest classifier was trained on the training set of HG-U133A and HG-U133A Plus 2.0, respectively, to prognosticate survival. Parameter was set at the upper limit of the total number of events in the training set to maintain equal sampling from patients who survived and those who experienced an event at 5 years. Tuning of random forest classifier parameters *mtry* (values 1, 2, 4, 6, 8, 10, 12, 14, 16, 18, 20, 22, 24) and *n*_*tree*_ (values 500, 1000, 2000, 5000) was done using grid. The best tuning parameters for the final classifier were selected based on the performance measure accuracy, as specified below.

The test dataset was evaluated using each of the tuned models to produce 0 or 1 to predict whether each patient died by 5 years. To calculate performance, patients alive at 5 years were considered to be true negatives (TNs) if the classifier correctly assigned them to good prognosis group, whereas they were considered as false negatives (FNs) if they died within 5 years. Similarly, patients who died within 5 years were considered to be true positives (TPs) if the classifier correctly assigned them to poor prognosis group, whereas they were considered as false positives (FPs) if they were alive at 5 years. Subsequently, sensitivity, specificity, and accuracy were calculated accordingly. The area under the receiver operator curve (AUC) was calculated based on the receiver operator characteristic (ROC) analysis using the random forest classification probability (pROC v1.8). The final tuning parameters selected were those that yielded the highest accuracy.

### Ensemble classifier: engineered variables

Following random forest classification using only the risk classification votes from different pipeline variants, classifiers were constructed using summary statistics as additional features. The engineered summary variables capture the total number of poor prognosis votes based on the variable aspects of preprocessing pipelines as follows: total number votes overall, total number of votes for pipelines using separate preprocessing, total number of votes for pipelines using merged preprocessing, total number of votes for RMA pipelines, total number of votes for GCRMA pipelines, total number of votes for MBEI pipelines, total number of votes for MAS5 pipelines, total number of votes for log_2_ MBEI pipelines, total number of votes for log_2_ MAS5 pipelines, total number of votes for RMA and MAS5 pipelines, total number of votes for pipelines using default annotation, and total number of votes for pipelines using alternative annotation. The derivation of engineered variables is summarized in Additional file Table S4.

Random forest models were built upon the following feature combinations: ensemble of preprocessing pipeline variants and the engineered variables, ensemble of engineered variables, and ensemble of only feature variables selected by the Boruta algorithm (Boruta v4.0.0). Random forest models were tuned based on performance similar to above. For the HG-U133A dataset, models were constructed by incorporating all patients in the cohort or only the subset of patients with unanimous agreement across the preprocessing pipelines. For the HG-U133A Plus 2.0 dataset, given the smaller sample size, models were constructed by incorporating all patients in the cohort to maintain sufficient statistical power.

### Classifier evaluation

The prognostic performance of the tuned classifiers was evaluated on the test set Kaplan-Meier estimates with the log-rank test and unadjusted Cox proportional hazard ratio model used to compare between the two groups (survival v2.38.0). In order to assess the performance of random forest-based ensemble classifiers, we compared the random forest classifier hazard ratio (HR), the HR in the subset of patients with unanimous agreement across 24 preprocessing pipelines, as well as the HR of individual preprocessing pipelines. Similarly, binary classification measure accuracy was compared. To compare between the random forest classifiers derived for each hypoxia signature, we assessed prognostic performance using the AUC. The ROC analysis was conducted for each signature using the random forest classification probability (pROC v1.8).

### Statistical comparison analysis

We compared the HR performance in the array of random forest classifier models for each hypoxia signature. The classifier HRs were split based on the features used to build the classifier: preprocessing pipelines, engineered variables, and feature variable selection. A paired t-test was used to assess statistical differences in the log_2_-transformed Hazard Ratios.

### New signature creation using preprocessing ensembles

Using the HG-U133A platform datasets, we sought to elucidate the ability of preprocessing ensemble classifiers to improve upon performance of novel signatures. To this end, we generated a 100-top-ranked-gene novel signature for individual preprocessing pipelines. This was done for preprocessing pipelines where all HG-U133A datasets were preprocessed together, yielding 12 individual signatures. To ascertain the signatures, each preprocessing normalization method was used to median-dichotomize the patient cohort by low or high abundance for each gene. The unadjusted Cox proportional hazard model was used to determine the univariate performance of individual genes to prognostic outcome. Statistical significance was assessed using the Wald test and *p*-values were false-discovery rate (FDR) adjusted to correct for multiple-testing. The 100 top-ranked genes with adjusted *p*-values < 0.05 were selected to constitute the signature. The individual signatures from the 12 preprocessing pipelines were validated using random forest classifiers using 10-fold cross-validation, where the random forest classifiers were trained on a training set and internally validated on a separate test set. The 12 good versus poor prognosis classifications were subsequently combined in a meta-random forest to evaluate its ability to predict prognosis compared to individual signature classifiers. The random forest model parameters were tuned as described above.

### Program usage

All statistical analyses and plotting were performed in R statistical environment (v3.2.1). The following packages were used for statistical analyses: randomForest v4.6.10, Boruta v4.0.0, survival v2.38.0, and pROC v1.8. All plots were generated in R using custom scripts for lattice (v0.2.31) and latticeExtra (v0.6.26).

## Results

### Study design: ensembles of preprocessing pipelines

Our overall approach to evaluate non-linear preprocessing ensembles is outlined in Figure 1. Our goal was to determine how multiple pre-processing methods might best be combined to improve biomarkers predictive of patient prognosis. The datasets used were separated based on the microarray platform – HG-U133A and HG-U133 Plus 2.0 – because of previously reported differences in their noise characteristics [19]. The union of all HG-U133A datasets contains 1,564 patients while that of the HG-U133 Plus 2.0 datasets contains 579. Each individual dataset was preprocessed using 24 pipeline variants, and then each hypoxia signature was scored for each pre-processing variant. This resulted in 24 predictions for each combination of patient and signature. Additionally, we derived several engineered variables from counting the total number of votes based on various preprocessing pipeline characteristics (**Additional file 4: Table S4**). Random forest classifiers were constructed to predict prognosis for individual patients using combinations of the ensemble of 24 preprocessing pipeline predictions and the engineered features. We evaluated the performance of these classifiers using Kaplan-Meier analysis, Cox proportional hazard model, and the binary classification accuracy.

**Figure 1.**
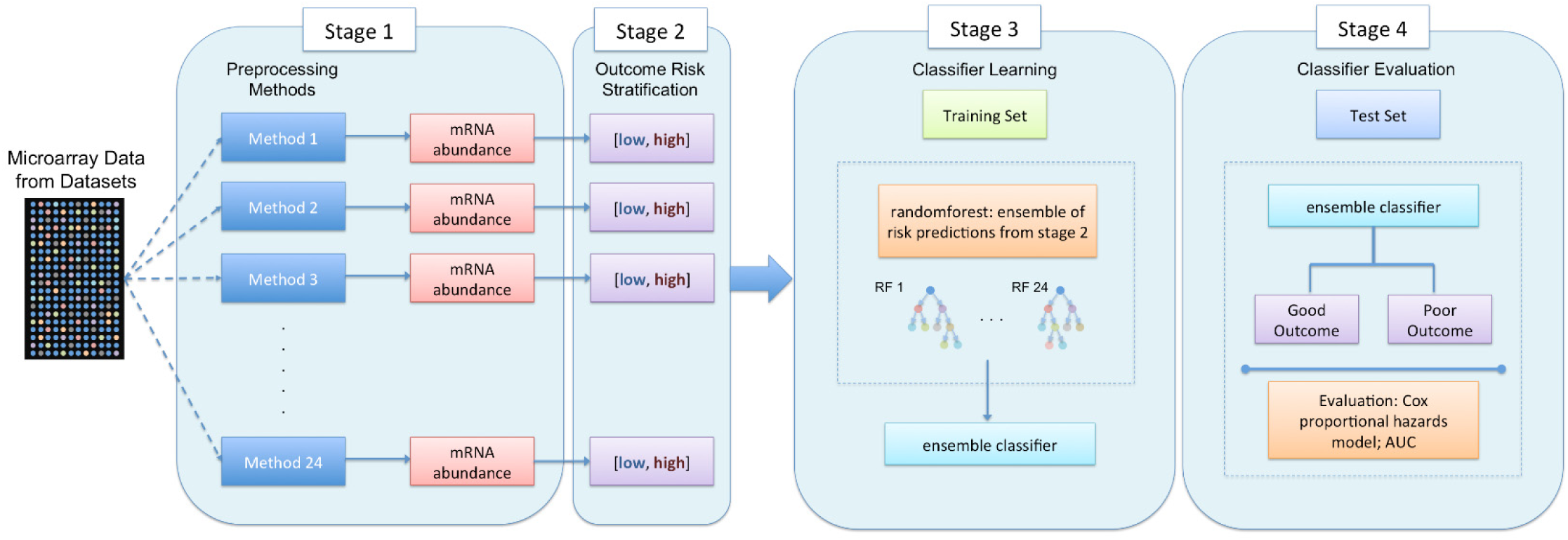
Summary of the study design for ensemble classification for evaluation of a biomarker. Microarray data are obtained from specific platforms and preprocessing using 24 different pipelines to normalize the mRNA gene expression. Risk groups are then assigned based on the biomarker of interest, resulting in a collection of either good or poor prognosis stratification based on the expression obtained from various preprocessing methods. Stratification into either good or poor prognosis represents a vote for that group, resulting in a score between 0 and 24. The ensemble of classifications is combined as features for random forest based machine learning. Random forest classifiers learning on a selected training set and evaluated on the test set. The robustness of the classifier derived for the biomarker of interest is evaluated with Cox proportional hazard ratio modeling and Kaplan-Meier survival estimates.

### Different preprocessing ensembles perform best for different biomarkers

We compared the performance of the individual preprocessing pipelines with to those of ensemble approaches. This process was conducted for each of the four hypoxia signatures and both microarray platforms. **Additional file 5 and 6: Table S5** (HG-U133A) and **S6** (HG-U133 Plus 2.0) comprise the hazard ratios (HRs) and 95% confidence intervals (CIs) determined for each of the 24 preprocessing pipelines, the random forest classifiers evaluated, and the simple preprocessing unanimous classifier, for each signature. Note that in this design each classifier is evaluated on a fully-independent validation cohort, to mitigate over-fitting.

Figure 2A shows a representative forest plot of the prognostic ability of various classifiers measured in HRs for the Winter metagene signature, using the HG-133A microarray platform. The best prediction of prognosis was observed in the subset of patients with unanimous agreement across the pipelines [HR 3.48, 95% confidence interval (CI) 2.44-4.95, *p* = 4.99 × 10^−12^]. However, the unanimous classification method only makes predictions for 41% (642) of patients while the remainder are unclassified. With incorporation of all patients in the HG-U133A dataset, the random forest classifier using engineered variables derived from votes of preprocessing pipeline features appeared to be a better predictor of prognosis than any individual pipelines (HR 2.39, 95% CI 1.94-2.93, *p* = 9.89 × 10^−17^). Similarly, the prognostic ability of two other ensemble random forest classifiers (preprocessing pipeline in combination with engineered variables, and preprocessing pipelines ensemble) also performed better than any individual pipelines (HR 2.25, 95% CI 1.83-2.76, *p* = 9.59 × 10^−15^ and HR 2.24, 95% CI 1.82-2.75, *p* = 1.41 × 10^−14^).

**Figure 2.**
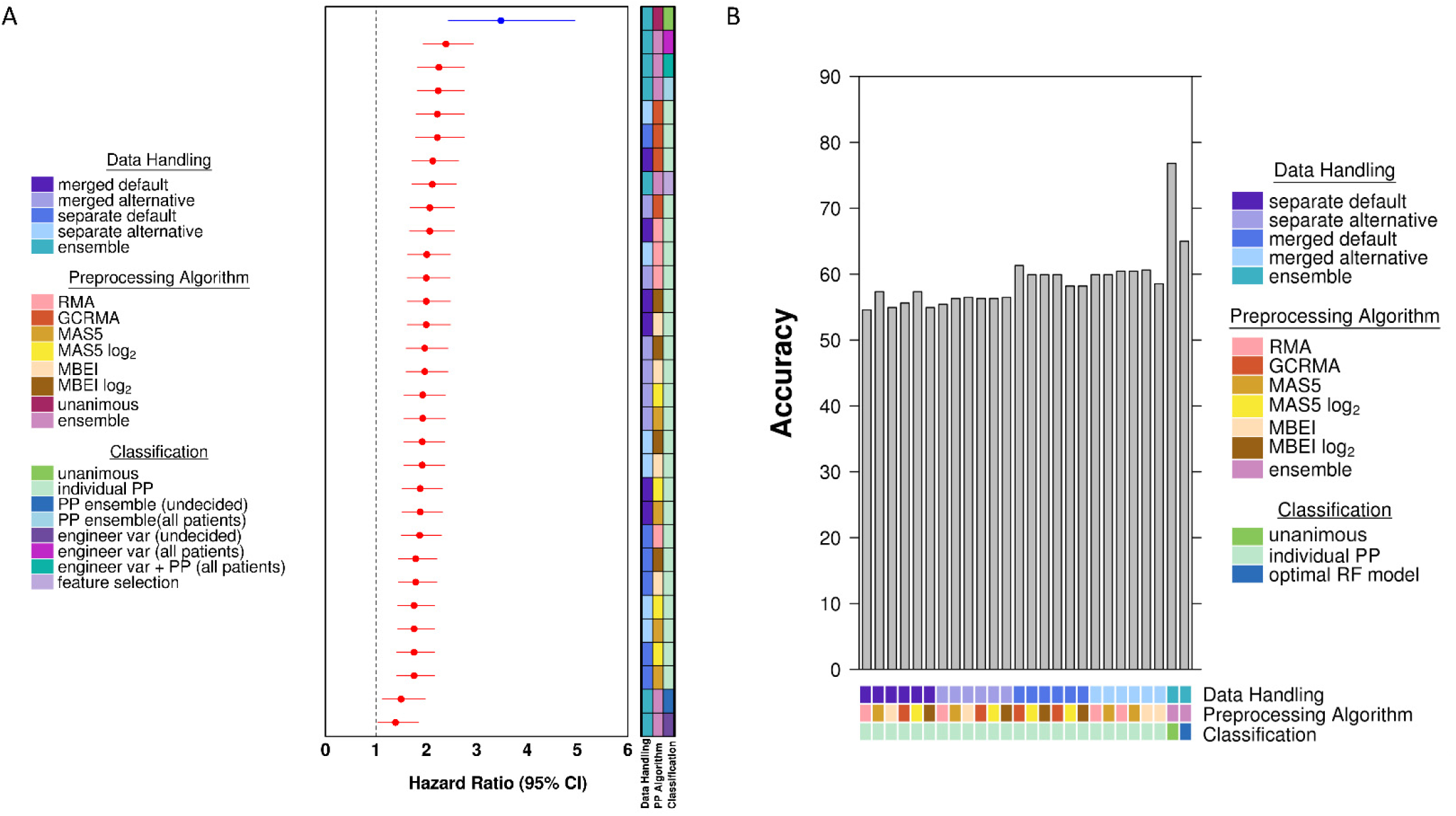
Representative hazard ratio forest plot and accuracy for Winter metagene signature using the HG-U133A microarray platform. (A) Forest plot of log_2_ hazard ratios with 95% confidence intervals obtained for each of the 24 preprocessing (PP) methods, the random forest classifiers evaluated, and the simple unanimous vote classifier (total number of votes for poor prognosis either 0 or 24). The forest plot is ordered as decreasing hazard ratio. The dotted line represents a hazard ratio of 1. The blue hazard ratio with its 95% confidence interval represents the hazard ratio for the simple unanimous vote classifier. (B) Bar plot of accuracy obtained for each of the 24 preprocessing methods, the random forest classifiers evaluated, and the simple unanimous vote classifier. The bars are ordered by preprocessing pipelines, the unanimous classifier, and the best performing random forest classifier, from left to right.

Surprisingly, though, this improved performance of random forest classifier of pre-processing methods was not a general feature of signatures. Rather, the performance of the ensemble classifier in comparison to individual pipeline variants was highly variable for the Buffa (**Additional file 8: Figure S1**) Hu (**Additional file 9: Figure S2**) and Sorensen signatures (**Additional file 10: Figure S3**). Further, the combination of features resulting in the best classifier was not consistent across the four signatures: engineered variables were important for the Buffa and Winter signatures (Buffa: HR 2.15, 95% CI 1.75-2.64, *p* = 4.03 × 10^−13^; Winter: HR 2.39, 95% CI 1.94-2.93, *p* = 9.89x 10^−17^), but feature selection using the Boruta algorithm yielded the highest performing classifier for Hu (HR 1.63, 95% CI 1.32-2.00, *p* = 3.87 × 10^−6^) and Sorensen signatures (HR 2.28, 95% CI 1.87-2.78, *p* = 2.51 × 10^−16^).

These findings of strong divergence in the best way to merge pre-processing algorithms held when we considered other metrics of classification accuracy besides HRs. For example, classification accuracy and evaluation of the area under the receiver operating characteristics curve (AUC) again show the benefits of specific pre-processing ensembles for the Winter signature (Figure 2B) matching those in the HR analysis, and analogously for the Buffa (**Additional file 8: Figure S1**), Hu (**Additional file 9: Figure S2**), and Sorensen signatures (**Additional file 10: Figures S3**).

These trends were also independent of the specific microarray platform used: results were comparable in patients analyzed using the HG-U133 Plus 2.0 microarray platform (**Additional file 11-14: Figure S4-S7**). The preprocessing unanimous classifier based on simple risk voting resulted in superior prognostication compared to individual preprocessing variants for all signatures except for Sorensen. Furthermore, the random forest classifiers evaluated did not improve upon unanimous classification, except for the Sorensen signature. The best performing random forest classifier was also inconsistent and variable across the biomarkers evaluated. The Kaplan-Meier plots for the HG-U133A dataset are shown in **Additional file 15-18: Figure S8-S11**. The Kaplan-Meier plots for the HG-U133 Plus 2.0 dataset are shown in **Additional file 19-22: Figure S12-S15**.

### Comparison of patient prognosis prediction between signature classifiers

Taken together, our results show that it is possible to improve upon individual pre-processing pipelines using ensemble techniques, but that the best way to assemble these ensembles varies with the biomarker signature, and not the microarray platform. Figure 3A compares the best ensemble of pre-processing methods to the best individual preprocessing method for each signature and microarray platform. Consistent with our previous results, the random forest classifier outperformed the preprocessing method for Winter and Sorensen signatures, but not for Buffa and Hu signatures. The ROC curve and corresponding AUC obtained for the best ensemble of preprocessing strategies is shown in Figure 3B and Figure 3C. The Buffa, Winter, and Sorensen signature classifiers demonstrated similar AUCs for mortality risk stratification between the two microarray platforms. Conversely, the Hu signature classifier showed better risk stratification using the HG-U133 Plus 2.0 platform compared to the HG-U133A platform.

**Figure 3.**
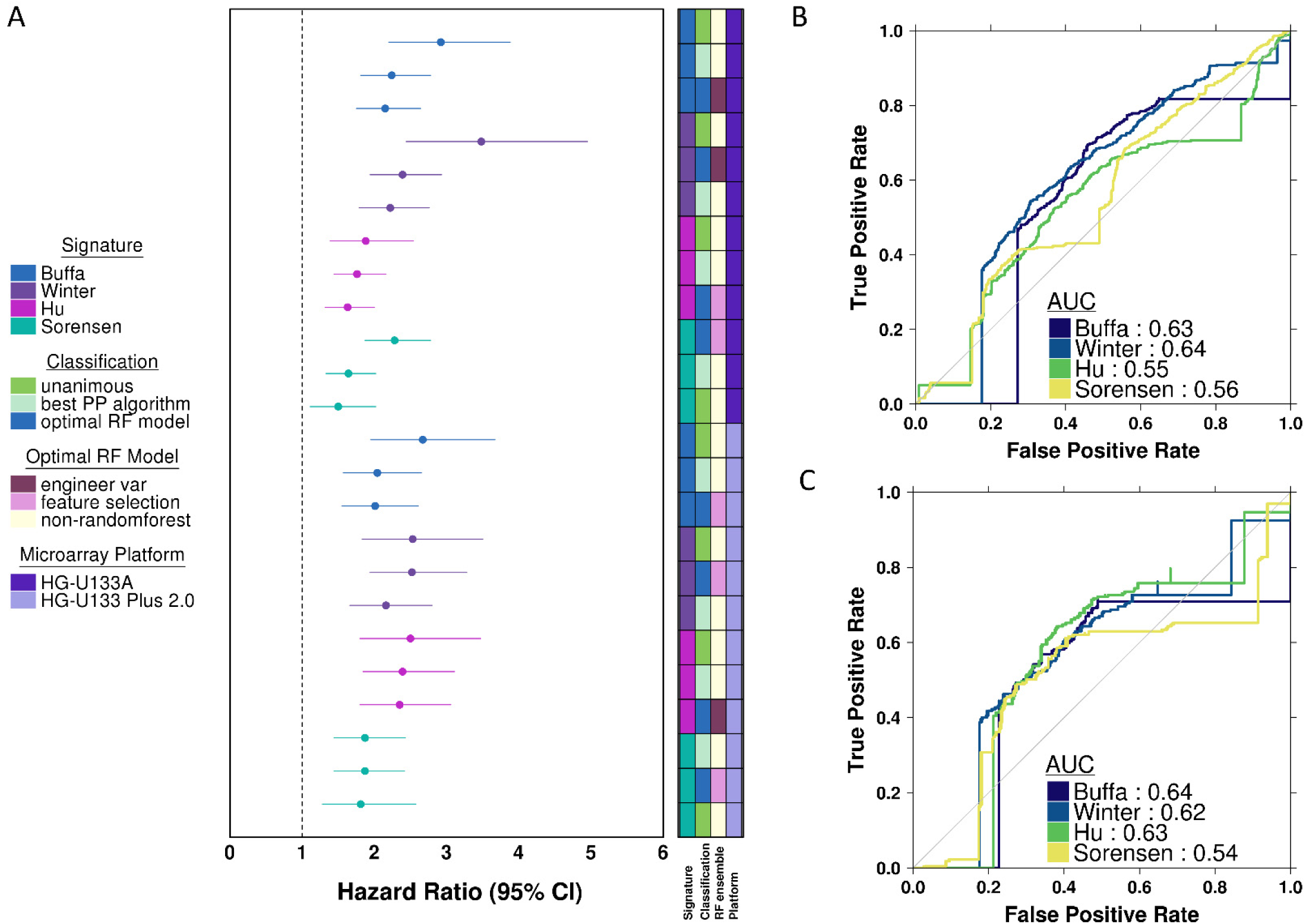
Summary hazard ratio forest plot and receiver operator curves. (A) Forest plot of log_2_ hazard ratios with 95% confidence intervals obtained for the best performing preprocessing method, best performing random forest classifier, and the unanimous vote classifier. Plot is ordered by decreasing hazard ratio within each signature and microarray platform evaluated. Colors correspond to the specific signature evaluated. (B and C) Receiver operator curves and area under the curve (AUC) obtained from the best performing random forest classifier for each biomarker, as determined by the highest hazard ratio. HG-U133A ROC curves shown in A, and HG-U133 Plus 2.0 ROC curves shown in B.

To determine if there are general properties of an ensemble of preprocessing methods that contribute to its performance, we compared each classifier feature to the ultimate performance of the classifier. This was done separately for both microarray platforms. For the HG-U133A platform, patients where all preprocessing methods gave a consistent results (unanimous preprocessing agreement) were statistically easier to classify than those where there was divergence amongst the pre-processing methods. These patients are thus more difficult to prognose, even though ensembles do improve upon the best individual pre-processing method. Similarly for the HG-U133 Plus 2.0 platform, patients with unanimous preprocessing agreement were statistically significantly or trend significantly easier to classify than those with divergence across classifiers. This trend was consistent across all four signatures evaluated, and across both platforms, suggesting that there is a patient sub-group that is fundamentally easier to classify, and that on the agreement of pre-processing methods on this sub-group can give increased confidence to the accuracy of molecular biomarkers.

### Generalization to Non-Hypoxia Signatures

To assess the generality of these observations, we trained independent prognostic signatures on each pre-processing method (Figure 4). Thus the same training dataset was pre-processed in 12 distinct ways, and then a learner was applied to each of these, leading to 12 distinct prognostic biomarkers. We focused on the HG-U133A data for this experiment, given its larger sample-size. We selected a standard straight-forward machine-learning approach, involving feature-selection with a univariate statistical text (Cox proportional hazards modeling) and modeling using the non-metric random forest approach. We then evaluated whether these 12 separate classifiers gave similar predictions for each individual patient, and attempted to create an ensemble of them. Finally the twelve separate and one ensemble classifiers were validated on the independent validation dataset using the AUC and Cox proportional hazards modeling.

**Figure 4.**
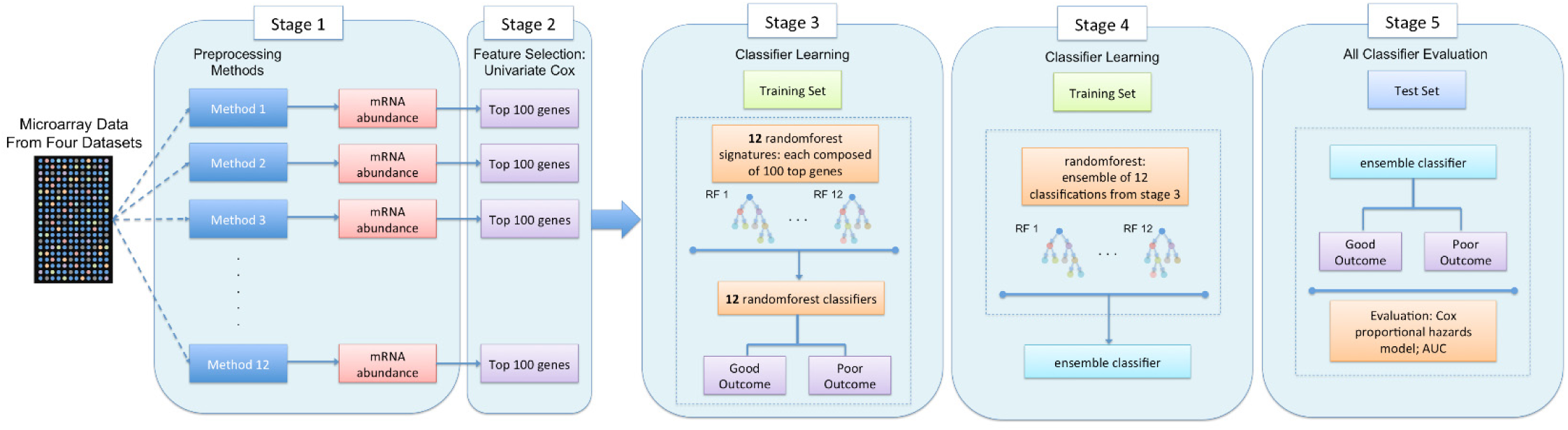
Summary of the study design for development of novel signature classifiers for each preprocessing pipeline and evaluating its performance in a meta-ensemble classifier. Microarray data are obtained from specific platforms and preprocessing using 24 different pipelines to normalize the mRNA gene expression. The gene expression is median dichotomized into two expression groups. Novel signatures are determined as the top 100 genes that reached significant after adjustment for false discovery rate, for each preprocessing pipeline (total 12). The training of a random forest classifier based on the individual novel signatures result in individual risk classifications of survival prognosis. These risk stratification are subsequently combined in a meta-random forest classifier to evaluate the robustness of the signature with Cox proportional hazard ratio modeling and Kaplan-Meier survival estimates.

The signatures trained with each of the 12 preprocessing pipelines had remarkably similar accuracy and HRs (Figure 5A), and a subset of genes overlapped across multiple signatures (**Additional file 7: Table S7**). An ensemble of these 12 classifiers resulted in marginally, but not statistically significant, improved predictions, suggesting that the signatures are not providing complementary information. To verify this, we compared the agreement of the per-patient predictions across all signatures. Figure 5B illustrates the predictions of individual signature classifiers across all patients stratified by the true survival outcome. The signature showed highly concordant classification, with patients with mortality events were similarly classified as having poor prognosis across the signatures and patients with continued survival were similarly classified as having good prognosis across the signatures. Similarly, inaccurate predictions of survival and mortality occurred in a comparable subset of patients across the signatures. Taken together, it appears that all signatures predict either good or poor survival for a similar cohort of patients, and that there remains a group of patients whose prognosis is difficult to predict and that leveraging orthogonal information from multiple pre-processing schemes will not help in making more accurate predictions for these.

**Figure 5.**
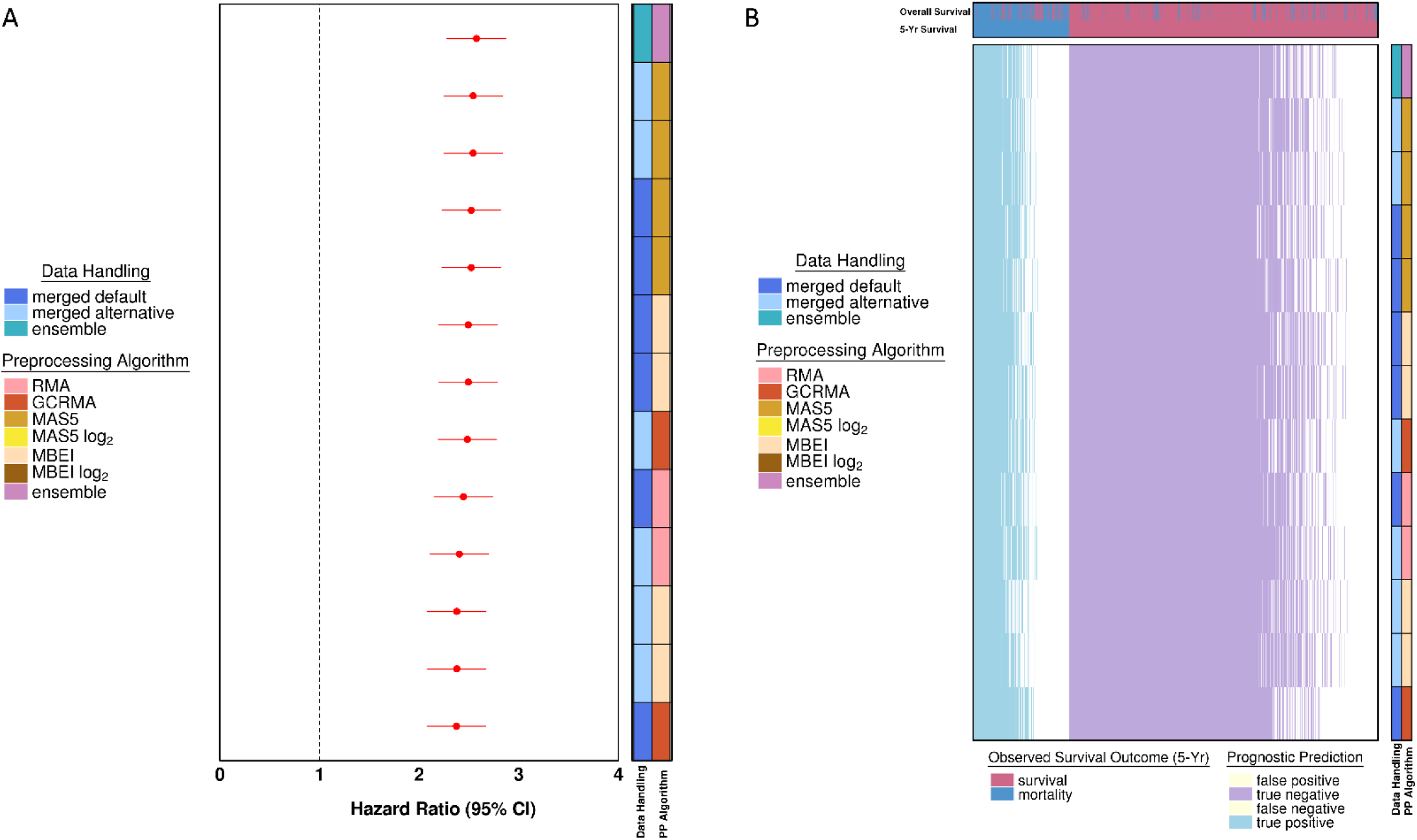
Hazard ratio forest plots of classifier performance and heatmap of individual classifier predictions of survival prognosis. (A) Forest plot with 95% confidence intervals of novel signature classifiers. The forest plot is ordered as decreasing hazard ratio. The dotted line represents a hazard ratio of 1. (B) Heatmap of classifier predictions of 5-year survival status. The classifiers (by row) from signatures are ordered by decreasing performance of patient outcome prediction. Patients (by column) are ordered by the degree of agreement of predictions across the array of novel signatures identified from 12 different preprocessing variant pipelines. The true outcome of patients is shown as either 5-year survival status or overall survival status up to the end of study follow-up. Blue represents true positives with correct prediction of poor prognosis. Purple represents true negatives with correct prediction of good prognosis. The white part of the heatmap represents incorrect predictions of good or poor prognosis.

## Discussion

Some groups have suggested that different preprocessing methods have minor effect on predictive signatures [41, 42]. Other work has suggested that this is incorrect, and that different preprocessing algorithms results in substantial differences in outcomes [18, 19]. Indeed we previously showed that ensemble classification combining preprocessing techniques using a unanimous voting method could identify high-confidence predictions, thereby giving increased confidence to risk stratification tools. We sought here to extend this approach and to discover if the predictions from multiple pre-processing algorithms might be combined into more accurate ensemble calls.

Our results demonstrate that there is indeed value to leveraging multiple pre-processing techniques. However, they yield the surprising result that the optimal way to do so is dependent on the characteristics of an individual signature. That is, one must consider all pre-processing methodologies for each new biomarker to determine if and to what extent combining them will improve predictions: there is no apparent universal approach to optimize this problem, even holding the dataset constant. Further, in ensembles appear to be limited in the extent to which they can improve signatures – there remains a subset of hard-to-classify patients for whom varying characteristics of the pre-processing do not help in classification. Large inter-individual differences exist in a plethora of extrinsic factors that play an equally imperative role in driving survival outcomes. These include environmental exposure factors, socioeconomic factors, patient compliance concerns, patient preferences, and social habits [43]. Treatment factors include success of surgery such as extent of margins, factors involved in the delivery of adjuvant treatments, as well as variability in the decision-making process between the patient and the treating physician. Currently, much of this information is not considered in the evaluation of intrinsic biological pattern on prognosis. Optimal prediction of outcomes will likely necessitate the integration of both intrinsic and extrinsic information in the biomarker development process. These findings are thus highly consistent with that demonstrated by Tofigh *et al.*, whereby the prognosis for a subset of breast cancer patients was intrinsically more difficult to predict [44].

Our results are not without limitations. First, the datasets included in the analyses herein represent only therapy-naïve early breast cancer tumors. It is well known that cancer is a disease of many, given the inter-tumor and intra-tumor heterogeneity observed. This precludes generalizations of these results to other tumor types. Second, we used random forests to derive classifiers, but potentially other machine learning algorithms may yield different results. Third, our study focused on four previously published hypoxia signatures and it would be difficult to extrapolate our findings to other microarray-based signatures. Studies are needed to elucidate the findings herein for other clinically promising signatures. Lastly, we only used microarray datasets to assess the utility of random forest classifiers for risk stratification. It may be that preprocessing ensemble classifications will be of greater benefit in fields where existing preprocessing methods are less robust [45].

Taken together, our data further highlights the need to incorporate extrinsic factors not accounted for by intrinsic biological signals, in the pursuit of integrative signatures that will allow for the realization of precision oncology.

## Declarations

### Acknowledgements

The authors thank all members of the Boutros lab for helpful comments and suggestions. This study makes use of data generated by the Molecular Taxonomy of Breast Cancer International Consortium (METABRIC). Funding for the project was provided by Cancer Research UK and the British Columbia Cancer Agency Branch.

### Availability of data and material

All METABRIC data is publicly available [46].

### Authors’ contributions

Project conception: IYG, PCB. Statistical and bioinformatics analyses: IYG, NSF, PCB. Composed the first draft of the manuscript: IYG. All authors read and approved the final manuscript.

### Ethics approval and consent to participate

Not applicable.

## Additional Files

**Additional file 1:** Table S1. Overview of preprocessing algorithms.

**Additional file 2:** Table S2 Summary of 24 preprocessing methods.

**Additional file 3:** Table S3 Overview of hypoxia prognostic signatures.

**Additional file 4:** Table S4 Summary of votes used to calculate engineered variables.

**Additional file 5:** Table S5 Hazard ratios and 95% confidence intervals obtained for each of the 24 preprocessing methods, the random forest classifiers evaluated, and the simple unanimous vote classifier, per signature (HG-U133A microarray platform).

**Additional file 6:** Table S6 Hazard ratios and 95% confidence intervals obtained for each of the 24 preprocessing methods, the random forest classifiers evaluated, and the simple unanimous vote classifier, per signature (HG-U133 Plus 2.0 microarray platform).

**Additional file 7:** Table S7 Frequency of top ranked genes selected from each of the preprocessing pipelines by univariate Cox proportional hazard models.

**Additional file 8:** Figure S1 Hazard ratio forest plot and accuracy for Buffa metagene signature using the HG-U133A microarray platform. (A) Forest plot of log_2_ hazard ratios with 95% confidence intervals obtained for each of the 24 preprocessing (PP) methods, the random forest classifiers evaluated, and the simple unanimous vote classifier (total number of votes for poor prognosis either 0 or 24). The forest plot is ordered as decreasing hazard ratio. The dotted line represents a hazard ratio of 1. The blue hazard ratio with its 95% confidence interval represents the hazard ratio for the simple unanimous vote classifier. (B) Bar plot of accuracy obtained for each of the 24 preprocessing methods, the random forest classifiers evaluated, and the simple unanimous vote classifier. The bars are ordered by preprocessing pipelines, the unanimous classifier, and the best performing random forest classifier, from left to right.

**Additional file 9: Figure S2** Hazard ratio forest plot and accuracy for Hu signature using the HG-U133A microarray platform. (A) Forest plot of log_2_ hazard ratios with 95% confidence intervals obtained for each of the 24 preprocessing (PP) methods, the random forest classifiers evaluated, and the simple unanimous vote classifier (total number of votes for poor prognosis either 0 or 24). The forest plot is ordered as decreasing hazard ratio. The dotted line represents a hazard ratio of 1. The blue hazard ratio with its 95% confidence interval represents the hazard ratio for the simple unanimous vote classifier. (B) Bar plot of accuracy obtained for each of the 24 preprocessing methods, the random forest classifiers evaluated, and the simple unanimous vote classifier. The bars are ordered by preprocessing pipelines, the unanimous classifier, and the best performing random forest classifier, from left to right.

**Additional file 10**: **Figure S3** Hazard ratio forest plot and accuracy for Sorensen signature using the HG-U133A microarray platform. (A) Forest plot of log_2_ hazard ratios with 95% confidence intervals obtained for each of the 24 preprocessing (PP) methods, the random forest classifiers evaluated, and the simple unanimous vote classifier (total number of votes for poor prognosis either 0 or 24). The forest plot is ordered as decreasing hazard ratio. The dotted line represents a hazard ratio of 1. The blue hazard ratio with its 95% confidence interval represents the hazard ratio for the simple unanimous vote classifier. (B) Bar plot of accuracy obtained for each of the 24 preprocessing methods, the random forest classifiers evaluated, and the simple unanimous vote classifier. The bars are ordered by preprocessing pipelines, the unanimous classifier, and the best performing random forest classifier, from left to right.

**Additional file 11: Figure S4** Hazard ratio forest plot and accuracy for Buffa metagene signature using the HG-U133 Plus 2.0 microarray platform. (A) Forest plot of log_2_ hazard ratios with 95% confidence intervals obtained for each of the 24 preprocessing (PP) methods, the random forest classifiers evaluated, and the simple unanimous vote classifier (total number of votes for poor prognosis either 0 or 24). The forest plot is ordered as decreasing hazard ratio. The dotted line represents a hazard ratio of 1. The blue hazard ratio with its 95% confidence interval represents the hazard ratio for the simple unanimous vote classifier. (B) Bar plot of accuracy obtained for each of the 24 preprocessing methods, the random forest classifiers evaluated, and the simple unanimous vote classifier. The bars are ordered by preprocessing pipelines, the unanimous classifier, and the best performing random forest classifier, from left to right.

**Additional file 12: Figure S5** Hazard ratio forest plot and accuracy for Winter metagene signature using the HG-U133 Plus 2.0 microarray platform. (A) Forest plot of log_2_ hazard ratios with 95% confidence intervals obtained for each of the 24 preprocessing (PP) methods, the random forest classifiers evaluated, and the simple unanimous vote classifier (total number of votes for poor prognosis either 0 or 24). The forest plot is ordered as decreasing hazard ratio. The dotted line represents a hazard ratio of 1. The blue hazard ratio with its 95% confidence interval represents the hazard ratio for the simple unanimous vote classifier. (B) Bar plot of accuracy obtained for each of the 24 preprocessing methods, the random forest classifiers evaluated, and the simple unanimous vote classifier. The bars are ordered by preprocessing pipelines, the unanimous classifier, and the best performing random forest classifier, from left to right.

**Additional file 13: Figure S6** Hazard ratio forest plot and accuracy for Hu signature using the HG-U133 Plus 2.0 microarray platform. (A) Forest plot of log_2_ hazard ratios with 95% confidence intervals obtained for each of the 24 preprocessing (PP) methods, the random forest classifiers evaluated, and the simple unanimous vote classifier (total number of votes for poor prognosis either 0 or 24). The forest plot is ordered as decreasing hazard ratio. The dotted line represents a hazard ratio of 1. The blue hazard ratio with its 95% confidence interval represents the hazard ratio for the simple unanimous vote classifier. (B) Bar plot of accuracy obtained for each of the 24 preprocessing methods, the random forest classifiers evaluated, and the simple unanimous vote classifier. The bars are ordered by preprocessing pipelines, the unanimous classifier, and the best performing random forest classifier, from left to right.

**Additional file 14: Figure S7** Hazard ratio forest plot and accuracy for Sorensen signature using the HG-U133 Plus 2.0 microarray platform. (A) Forest plot of log_2_ hazard ratios with 95% confidence intervals obtained for each of the 24 preprocessing (PP) methods, the random forest classifiers evaluated, and the simple unanimous vote classifier (total number of votes for poor prognosis either 0 or 24). The forest plot is ordered as decreasing hazard ratio. The dotted line represents a hazard ratio of 1. The blue hazard ratio with its 95% confidence interval represents the hazard ratio for the simple unanimous vote classifier. (B) Bar plot of accuracy obtained for each of the 24 preprocessing methods, the random forest classifiers evaluated, and the simple unanimous vote classifier. The bars are ordered by preprocessing pipelines, the unanimous classifier, and the best performing random forest classifier, from left to right.

**Additional file 15: Figure S8** Kaplan-Meier survival curves evaluating the prognostic ability of the Buffa metagene signature using HG-U133A microarray platform. (A) Prognostic ability of signature in patients with unanimous ensemble agreement across preprocessing pipelines. (B) Prognostic ability of signature classification using the best performing preprocessing pipeline. (C) Prognostic ability of signature classification using the best performing random forest-based ensemble of preprocessing pipelines. Hazard ratios and p-values are from Cox proportional hazard ratio modeling.

**Additional file 16: Figure S9** Kaplan-Meier survival curves evaluating the prognostic ability of the Winter metagene signature using HG-U133A microarray platform. (A) Prognostic ability of signature in patients with unanimous ensemble agreement across preprocessing pipelines. (B) Prognostic ability of signature classification using the best performing preprocessing pipeline. (C) Prognostic ability of signature classification using the best performing random forest-based ensemble of preprocessing pipelines. Hazard ratios and p-values are from Cox proportional hazard ratio modeling.

**Additional file 17: Figure S10** Kaplan-Meier survival curves evaluating the prognostic ability of the Hu signature using HG-U133A microarray platform. (A) Prognostic ability of signature in patients with unanimous ensemble agreement across preprocessing pipelines. (B) Prognostic ability of signature classification using the best performing preprocessing pipeline. (C) Prognostic ability of signature classification using the best performing random forest-based ensemble of preprocessing pipelines. Hazard ratios and p-values are from Cox proportional hazard ratio modeling.

**Additional file 18: Figure S11** Kaplan-Meier survival curves evaluating the prognostic ability of the Sorensen signature using HG-U133A microarray platform. (A) Prognostic ability of signature in patients with unanimous ensemble agreement across preprocessing pipelines. (B) Prognostic ability of signature classification using the best performing preprocessing pipeline. (C) Prognostic ability of signature classification using the best performing random forest-based ensemble of preprocessing pipelines. Hazard ratios and p-values are from Cox proportional hazard ratio modeling.

**Additional file 19: Figure S12** Kaplan-Meier survival curves evaluating the prognostic ability of the Buffa metagene signature using HG-U133 Plus 2.0 microarray platform. (A) Prognostic ability of signature in patients with unanimous ensemble agreement across preprocessing pipelines. (B) Prognostic ability of signature classification using the best performing preprocessing pipeline. (C) Prognostic ability of signature classification using the best performing random forest-based ensemble of preprocessing pipelines. Hazard ratios and p-values are from Cox proportional hazard ratio modeling.

**Additional file 20: Figure S13** Kaplan-Meier survival curves evaluating the prognostic ability of the Winter metagene signature using HG-U133 Plus 2.0 microarray platform. (A) Prognostic ability of signature in patients with unanimous ensemble agreement across preprocessing pipelines. (B) Prognostic ability of signature classification using the best performing preprocessing pipeline. (C) Prognostic ability of signature classification using the best performing random forest-based ensemble of preprocessing pipelines. Hazard ratios and p-values are from Cox proportional hazard ratio modeling.

**Additional file 21: Figure S14** Kaplan-Meier survival curves evaluating the prognostic ability of the Hu signature using HG-U133 Plus 2.0 microarray platform. (A) Prognostic ability of signature in patients with unanimous ensemble agreement across preprocessing pipelines. (B) Prognostic ability of signature classification using the best performing preprocessing pipeline. (C) Prognostic ability of signature classification using the best performing random forest-based ensemble of preprocessing pipelines. Hazard ratios and p-values are from Cox proportional hazard ratio modeling.

**Additional file 22: Figure S15** Kaplan-Meier survival curves evaluating the prognostic ability of the Sorensen signature using HG-U133 Plus 2.0 microarray platform. (A) Prognostic ability of signature in patients with unanimous ensemble agreement across preprocessing pipelines. (B) Prognostic ability of signature classification using the best performing preprocessing pipeline. (C) Prognostic ability of signature classification using the best performing random forest-based ensemble of preprocessing pipelines. Hazard ratios and p-values are from Cox proportional hazard ratio modeling.

